# PhenoSpD: an integrated toolkit for phenotypic correlation estimation and multiple testing correction using GWAS summary statistics

**DOI:** 10.1101/148627

**Authors:** Jie Zheng, Tom G. Richardson, Louise A. C. Millard, Gibran Hemani, Christopher Raistrick, Bjarni Vilhjalmsson, Philip Haycock, Tom R Gaunt

## Abstract

**Background:** Identifying phenotypic correlations between complex traits and diseases can provide useful etiological insights. Restricted access to individual-level phenotype data makes it difficult to estimate large-scale phenotypic correlation across the human phenome. State-of-the-art methods, metaCCA and LD score regression, provide an alternative approach to estimate phenotypic correlation using genome-wide association study (GWAS) summary statistics.

**Results:** Here, we present an integrated R toolkit, PhenoSpD, to 1) apply metaCCA (or LD score regression) to estimate phenotypic correlations using GWAS summary statistics; and 2) to utilize the estimated phenotypic correlations to inform correction of multiple testing for complex human traits using the spectral decomposition of matrices (SpD). The simulations suggest it is possible to estimate phenotypic correlation using samples with only a partial overlap, but as overlap decreases correlations will attenuate towards zero and multiple testing correction will be more stringent than in perfectly overlapping samples. In a case study, PhenoSpD using GWAS results suggested 324.4 independent tests among 452 metabolites, which is close to the 296 independent tests estimated using true phenotypic correlation. We further applied PhenoSpD to estimated 7,503 pair-wise phenotypic correlations among 123 metabolites using GWAS summary statistics from Kettunen et al. and PhenoSpD suggested 44.9 number of independent tests for theses metabolites.

**Conclusion:** PhenoSpD integrates existing methods and provides a simple and conservative way to reduce dimensionality for complex human traits using GWAS summary statistics, which is particularly valuable for post-GWAS analysis of complex molecular traits.

**Availability:** R code and documentation for PhenoSpD V1.0.0 is available online (https://github.com/MRCIEU/PhenoSpD).

## Introduction

Phenotypic correlations between complex human traits and diseases provide useful etiological insights. For GWAS meta-analysis, a lack of individual-level phenotype data makes it difficult to estimate the phenotypic correlation across human traits and diseases. Here we consider two methods that estimate phenotypic correlations using GWAS summary statistics: metaCCA (Cichonska et al., 2016) and bivariate LD score regression (Bulik-Sullivan et al., 2015b). The metaCCA framework estimates phenotypic correlation between two traits based on a Pearson correlation between two univariate regression coefficients (betas) across a set of genetic variants; The bivariate LD score regression approach estimates the phenotypic correlation amongst the overlapping samples of two GWAS.

The recently developed MR-Base (Hemani et al.2016) and LD Hub (Zheng et al., 2017) tools include harmonized GWAS summary-level results. This provides an opportunity to estimate the phenotypic correlation structure across a wide range of high-dimensional, complex molecular traits, such as metabolites, that are potentially highly correlated. Bonferroni correction would markedly overcorrect for the inflated false-positive rate in such correlated datasets, resulting in a reduction in power. An appropriate method to correct for multiple testing among human traits and diseases based on the spectral decomposition of matrices (SpD) (Nyholt, 2004; Li and Ji, 2005) is considered in this study.

## Methods

### Overview of PhenoSpD

Figure 1 illustrates the key steps of the proposed pipeline, PhenoSpD: step (1) harmonise GWAS summary results from the same sample; step (2) apply the harmonized GWAS results to metaCCA or LD score regression to estimate the phenotypic correlation matrix of the traits; step (3) apply the phenotypic correlation matrix to the SpD approach and estimate the number of independent variables among the traits.

**Figure 1.**
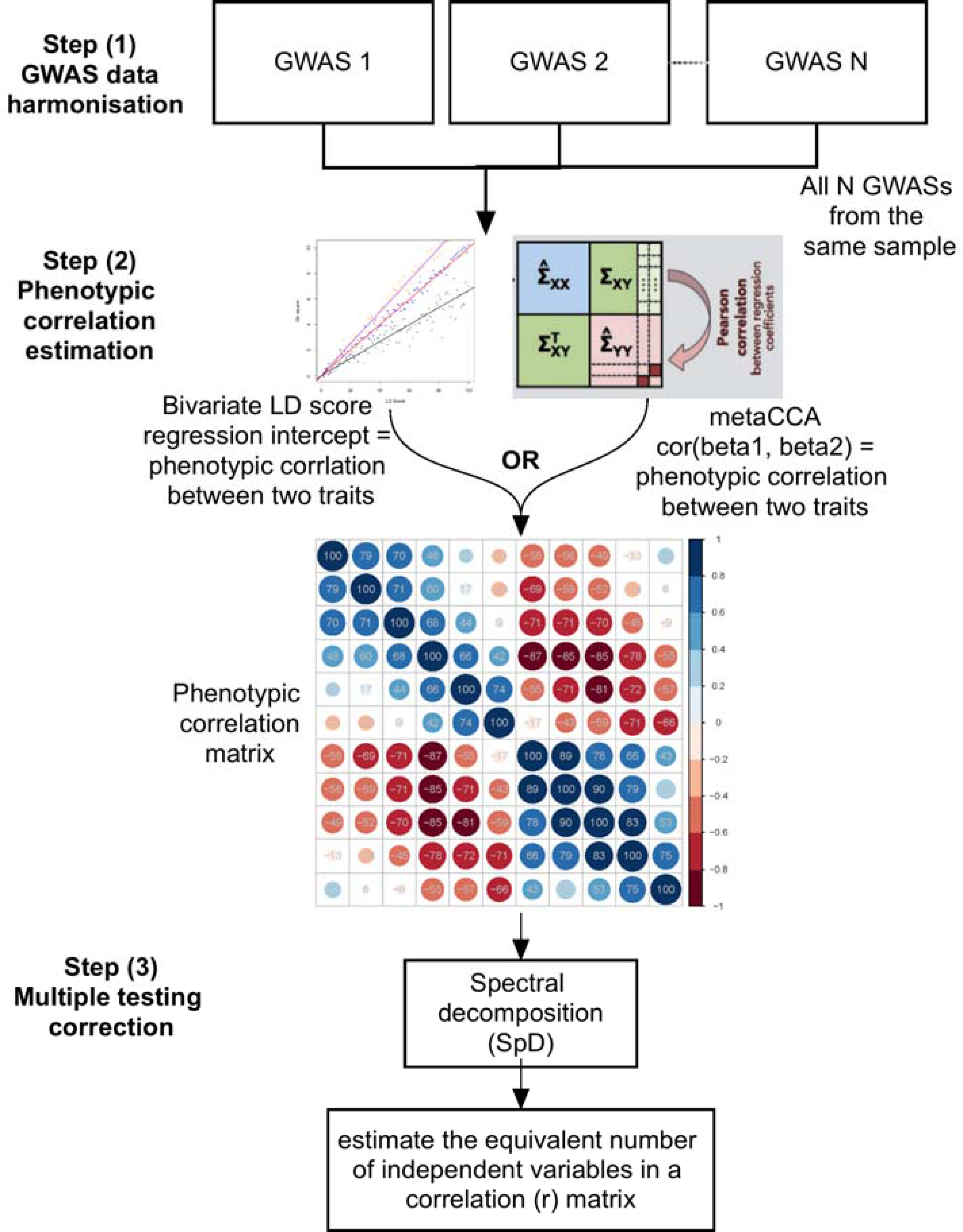
Flowchart of PhenoSpD.

### Simulation and validation of phenotypic correlation estimation

Firstly, we simulated the influence of the number of single nucleotide polymorphisms (SNPs), sample sizes of two GWASs and sample overlap between two GWASs on the accuracy of the phenotypic correlation estimation. As shown in Figure 2, we first created two samples A and B with different number of individuals (from 300 to 10,000 individuals), where the sample overlap between sample A and B ranged from 10% to 90%. Within each sample, we further assume complex human traits were influenced by both genetic and environmental factors. We simulated the phenotype data of two correlated human traits (phenotype 1 and phenotype 2) based on varying numbers of genetic factors (ranging from 10 to 10,000 SNPs) and 100 environmental factors. We then simulated the genotype data in each sample. After simulating the two phenotypic traits and the genotypic data in sample A and B, we then conducted four GWASs (GWASs of pheno-type 1 in sample A and B; GWASs of phenotype 2 in sample A and B) and recorded the summary statistics of these GWASs. To measure the accuracy of phenotypic correlation using GWAS summary statistics, we (1) calculated the observational phenotypic correlation (the Pearson correlation) of trait 1 and trait 2 in sample A and B separately; (2) estimated the phenotypic correlation of trait 1 and trait 2 using GWAS summary statistics of sample A and B separately. We simulated step (2) 100 times and estimate the mean and standard deviation of the estimated phenotypic correlations. Finally, we compared the estimated phenotypic correlation with the observational phenotypic correlation and recorded the deviation between observed and estimated correlations. To demonstrate the simulation systematically, we set up 4 groups of comparisons: (i) tested the influence of sample size; (ii) simulated the influence of sample overlap; (iii) estimated the influence of unbalanced sample size in sample A and B; (iv) tested the influence of number of SNPs. The R script for this simulation were provided as an supplementary file (simulation.R).

**Figure 2.**
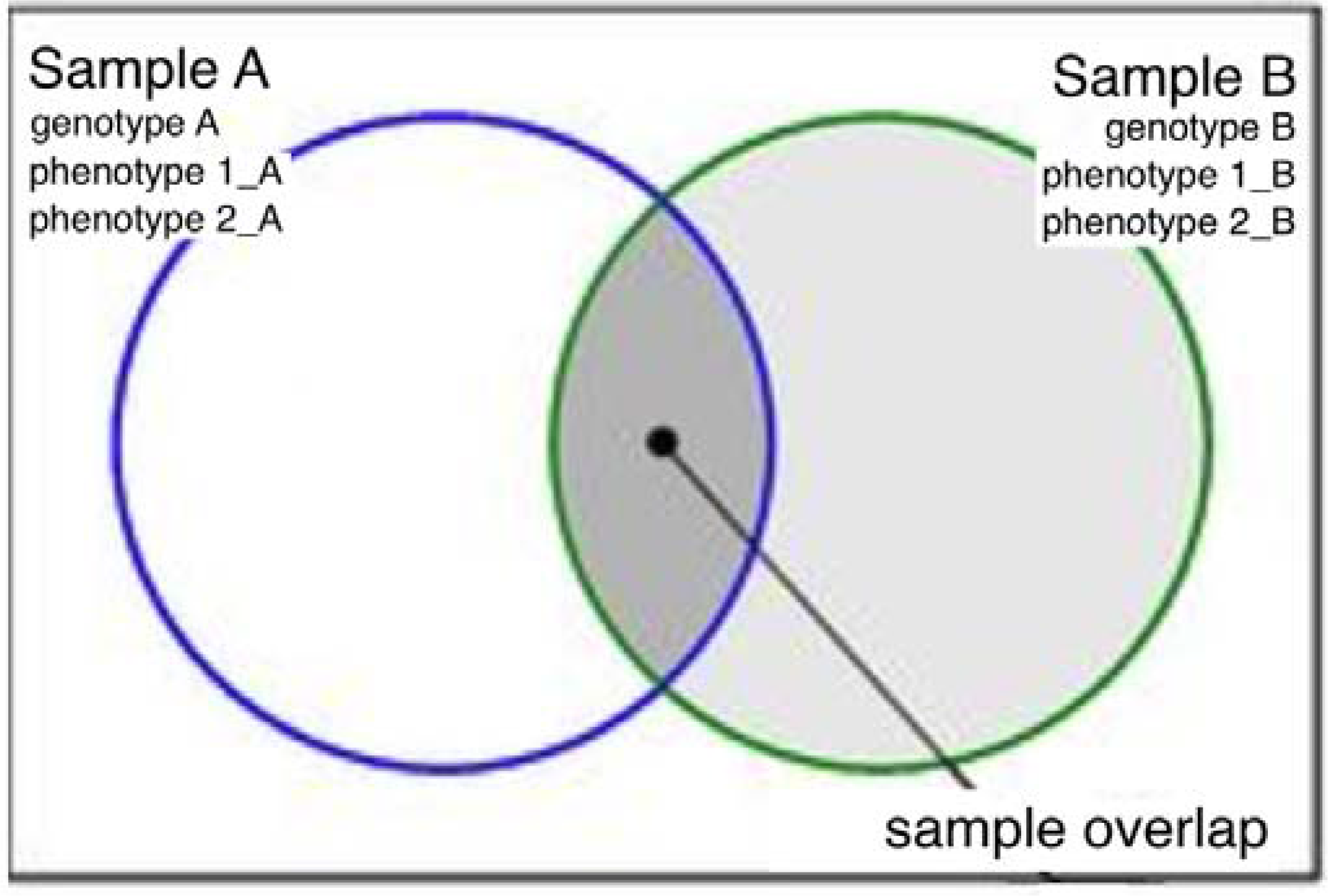
Demonstration of the simulation. For two samples A and B, we simulated the genotype data and phenotype data of two correlated human traits, phenotype 1 and phenotype 2. The sample overlap between sample A and B were ranged from 10% to 90% in this simulation.

We then tested the accuracy of phenotypic correlation estimation using GWAS summary statistics of 452 metabolites from Shin’s (Shin et al, 2014). Shin et al. also reported the observational phenotypic correlation in the supplementary table (Table S4), which was used as bench mark of our real case accuracy test.

Based on the simulation and real case validation, we listed our traits selection criteria in Table S1.

### Estimating the phenotypic correlations

Within our GWAS summary results database containing 1094 human traits, we selected 123 metabolites from Kettunen et al as a real case application (Kettunen et al, 2016) since these complex molecular traits are potentially highly correlated. We then applied metaCCA to these 123 metabolites to estimate the phenotypic correlation matrix (Table S2). Among the 123 metabolites, we further applied LD score regression to 107 of them (Table S3), which fit the assumptions of LD score regression (traits with large sample size (e.g. N > 5,000), good SNP coverage (e.g. number of SNPs > 200,000) and heritable (e.g. SNP heritability > 0.05)).

### Multiple testing correction for human traits

We applied the SpD approach to correct for multiple testing among the 123 metabolites. We implemented the R code of the well-known method, SNPSpD (Nyholt, 2004; Li and Ji, 2005), to estimate the number of independent traits using the phenotypic correlation matrix as input (Fig. 1). The output of the SpD function is the estimated the number of independent tests.

## Results

### Evaluation of phenotypic correlation estimation using simulated and real GWAS summary data

Table 1 show the influence of number of SNPs, sample sizes of two GWASs and sample overlap between two GWASs on the accuracy of the phenotypic correlation estimation. We found that the accuracy of phenotypic correlation estimation is mainly influenced by the number of overlapped individual in two GWAS studies. For example, the deviation between observed and estimated phenotypic correlation (Deviation_obs_est) improved from 82.1% to 5.8% when the percentage of sample overlap between two samples increased from 10% to 90%. Base on this simulation, we only applied metaCCA to GWASs from the same study to maximize the sample overlap between GWASs. In addition, we observed that the number of SNPs in GWAS will also influence the accuracy of the phenotypic correlation estimation. We should include as many SNPs as possible to maximize the accuracy of the estimation. Besides, we did not observe any major influence of the sample size of the GWAS on the accuracy of the estimation, so we included metabolites GWASs with various of sample sizes.

**Table 1.** The influence of number of SNPs, sample sizes of two GWASs and sample overlap between two GWASs on phenotypic correlation estimation.

| Model | N_ind_A | N_ind_B | N_overlap | Overlap_% | N_SNPs | N_EnvF | N_simu | y1_y2_A_obs | y1_y2_B_obs | Mean_y1_y2_est | SD_y1_y2_est | Deviation_obs_est (%) |
| --- | --- | --- | --- | --- | --- | --- | --- | --- | --- | --- | --- | --- |
| sample size 1 | 300 | 300 | 150 | 50% | 1000 | 100 | 100 | -0.70 | -0.70 | -0.46 | 0.56 | 34.1% |
| sample size 2 | 500 | 500 | 250 | 50% | 1000 | 100 | 100 | -0.71 | -0.70 | -0.47 | 0.56 | 33.0% |
| sample size 3 | 1000 | 1000 | 500 | 50% | 1000 | 100 | 100 | -0.70 | -0.70 | -0.47 | 0.54 | 33.3% |
| sample size 4 | 3000 | 3000 | 1500 | 50% | 1000 | 100 | 100 | -0.70 | -0.70 | -0.46 | 0.54 | 33.6% |
| sample size 5 | 5000 | 5000 | 2500 | 50% | 1000 | 100 | 100 | -0.70 | -0.70 | -0.47 | 0.54 | 33.2% |
| sample size 6 | 10000 | 10000 | 5000 | 50% | 1000 | 100 | 100 | -0.71 | -0.71 | -0.47 | 0.54 | 33.9% |
| sample overlap 1 | 5000 | 5000 | 1000 | 10% | 1000 | 100 | 100 | -0.70 | -0.70 | -0.13 | 0.39 | 82.1% |
| sample overlap 2 | 5000 | 5000 | 2000 | 20% | 1000 | 100 | 100 | -0.70 | -0.70 | -0.23 | 0.47 | 67.2% |
| sample overlap 3 | 5000 | 5000 | 3000 | 30% | 1000 | 100 | 100 | -0.71 | -0.71 | -0.33 | 0.47 | 54.0% |
| sample overlap 4 | 5000 | 5000 | 4000 | 40% | 1000 | 100 | 100 | -0.71 | -0.71 | -0.40 | 0.51 | 43.2% |
| sample overlap 5 | 5000 | 5000 | 5000 | 50% | 1000 | 100 | 100 | -0.71 | -0.71 | -0.47 | 0.53 | 33.3% |
| sample overlap 6 | 5000 | 5000 | 6000 | 60% | 1000 | 100 | 100 | -0.71 | -0.71 | -0.53 | 0.59 | 25.2% |
| sample overlap 7 | 5000 | 5000 | 7000 | 70% | 1000 | 100 | 100 | -0.70 | -0.70 | -0.58 | 0.57 | 17.5% |
| sample overlap 8 | 5000 | 5000 | 8000 | 80% | 1000 | 100 | 100 | -0.70 | -0.70 | -0.62 | 0.65 | 11.4% |
| sample overlap 9 | 5000 | 5000 | 9000 | 90% | 1000 | 100 | 100 | -0.71 | -0.71 | -0.67 | 0.67 | 5.8% |
| unbalance sample 1 | 5000 | 5000 | 9000 | 90% | 1000 | 100 | 100 | -0.71 | -0.71 | -0.67 | 0.67 | 5.8% |
| unbalance sample 2 | 5000 | 6000 | 9000 | 82% | 1000 | 100 | 100 | -0.71 | -0.71 | -0.64 | 0.66 | 9.8% |
| unbalance sample 3 | 5000 | 8000 | 9000 | 69% | 1000 | 100 | 100 | -0.70 | -0.70 | -0.58 | 0.65 | 17.3% |
| unbalance sample 4 | 5000 | 10000 | 9000 | 60% | 1000 | 100 | 100 | -0.70 | -0.70 | -0.54 | 0.62 | 22.9% |
| unbalance sample 5 | 5000 | 13000 | 9000 | 50% | 1000 | 100 | 100 | -0.70 | -0.70 | -0.48 | 0.54 | 30.9% |
| number of SNPs 1 | 5000 | 5000 | 2500 | 50% | 10 | 100 | 100 | -0.70 | -0.70 | -0.44 | 0.73 | 38.1% |
| number of SNPs 2 | 5000 | 5000 | 2500 | 50% | 100 | 100 | 100 | -0.70 | -0.70 | -0.48 | 0.53 | 34.1% |
| number of SNPs 3 | 5000 | 5000 | 2500 | 50% | 500 | 100 | 100 | -0.70 | -0.70 | -0.47 | 0.53 | 34.3% |
| number of SNPs 4 | 5000 | 5000 | 2500 | 50% | 1000 | 100 | 100 | -0.71 | -0.70 | -0.47 | 0.56 | 33.5% |
| number of SNPs 5 | 5000 | 5000 | 2500 | 50% | 5000 | 100 | 100 | -0.70 | -0.70 | -0.47 | 0.55 | 33.6% |
| number of SNPs 6 | 5000 | 5000 | 2500 | 50% | 10000 | 100 | 100 | -0.71 | -0.71 | -0.47 | 0.59 | 33.7% |

In this simulation, we compared the agreements of the observational (calculated from phenotypes) and estimated phenotypic correlation (estimated using GWAS results) of two human traits in two samples A and B. In general, we set up 4 groups of comparisons: (i) tested the influence of sample size; (ii) simulated the influence of sample overlap; (iii) estimated the influence of unbalanced sample size in sample A and B; (iv) tested the influence of number of SNPs. More details of the simulation can be found in methods section.
Abbreviations: N\_ind\_A and N\_ind\_B, number of individual in sample A and B. N\_overlap, number of overlapped samples in sample A and B. overlap\_%, the percentage of overlapped samples in A and B. N\_SNPs, number of SNPs in GWAS of sample A and B. N\_EnvF, number of environmental factors included in the model. N\_simu, number of simulations. y1\_y2\_A and y1\_y2\_B, the observed phenotypic correlation between two traits in sample A and B. Mean\_y1\_y2\_est, the mean value of the estimated phenotypic correlations in 100 simulations; SD\_y1\_y2\_est, the standard deviation of estimated phenotypic correlations in 100 simulations. Deviation\_obs\_est (%), the deviation between observational phenotypic correlation and estimated phenotypic correlation in each model of simulation.

We also tested the accuracy of phenotypic correlation estimation by comparing the observed phenotypic correlations (Table S4) and the estimated phenotypic correlation (Table S5) using real data from Shin’s (Shin et al, 2014). Figure 3 showed the estimated phenotypic correlations have a very high agreement with the observed phenotypic correlations (r^2^=0.89). The only exception is that some metabolites with large observed correlation have estimated correlation towards null.

**Figure 3.**
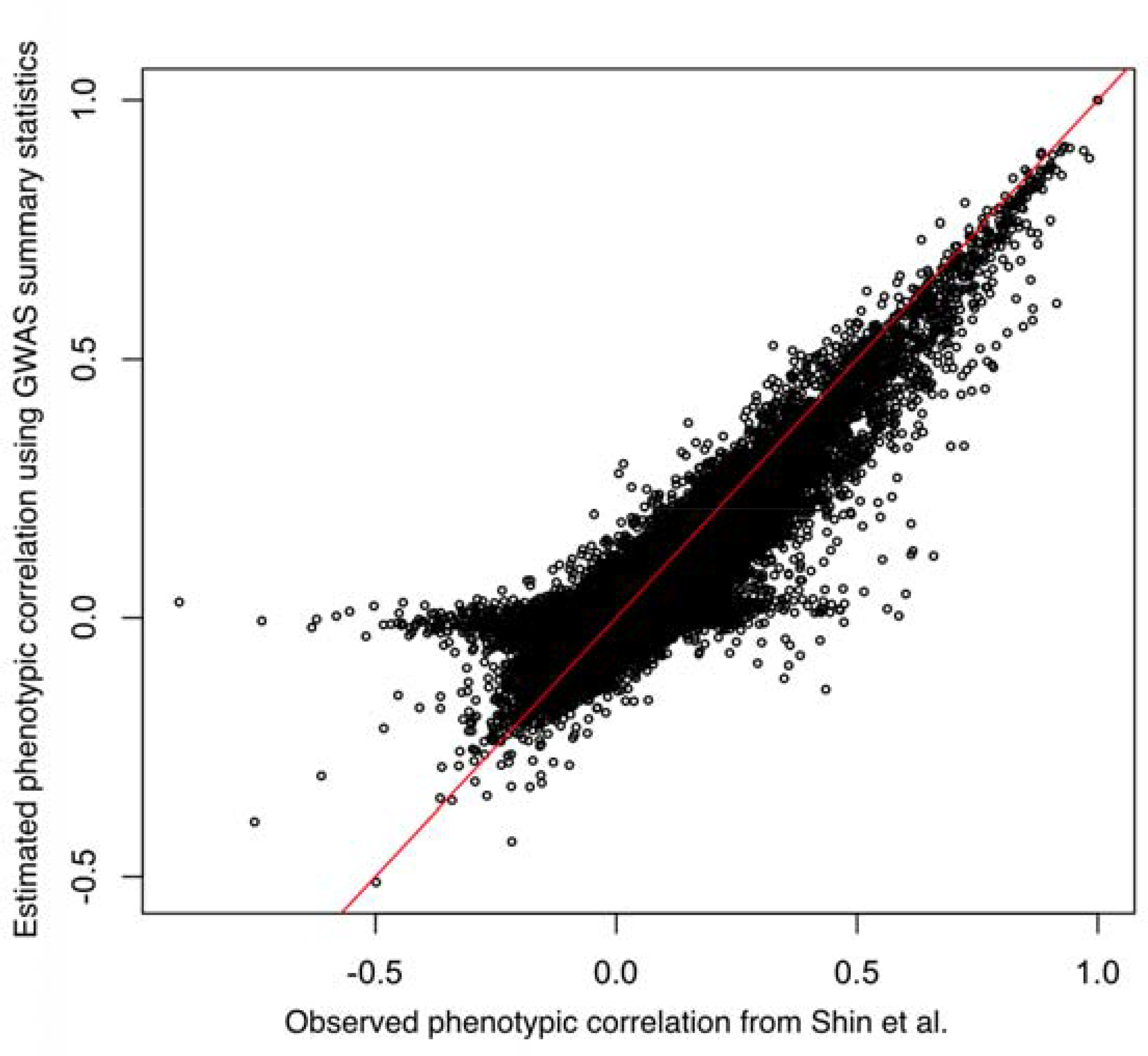
The comparison between the observed and estimated phenotypic correlations amongst 452 metabolites from Shin et al. Each point is one metabolite. The red line is X=Y line.

To further investigate this exception, we measured the difference between observed and estimated phenotypic correlations by using the sum of errors of phenotypic correlation (the sum of the differences between one metabolites on the rest metabolites). As show in Figure 4, the points with high level of errors (disagreements) are metabolites with limited sample size. The metabolites with limited sample size also have a limited number of sample overlap with other metabolites, which drive the phenotypic correlation estimation towards null.

**Figure 4.**
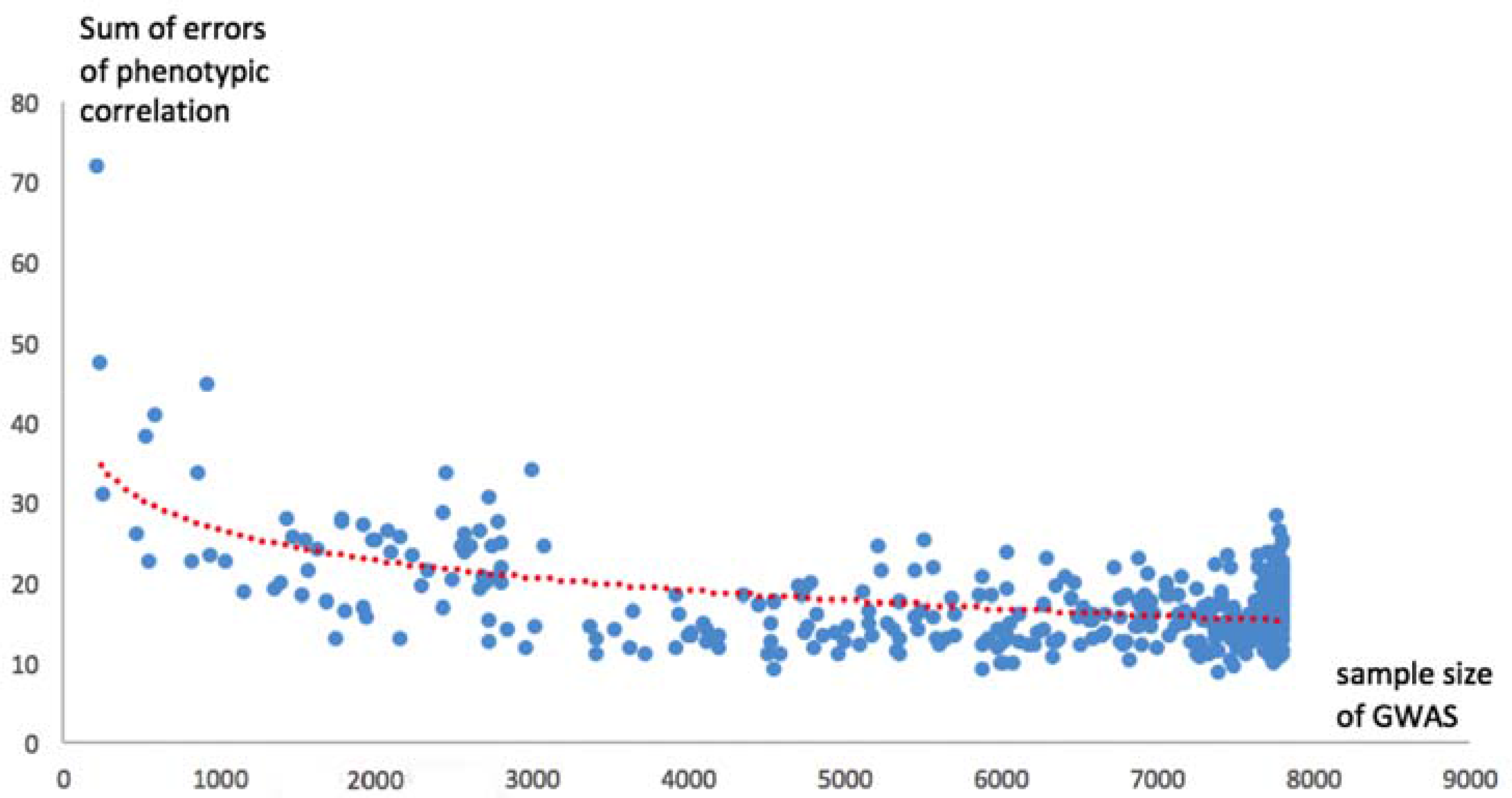
The relationship between sample size (sample overlap) and errors in 452 metabolites from Shin et al. Each point in the scatter plot is one metabolite. The x-axis is the sample size of each metabolite GWAS. The sum of errors of phenotypic correlation (y-axis) is refer to the difference between observed and estimated phenotypic correlations of one metabolite on the rest of the metabolites.

### A practical comparison between metaCCA and LD score regression on estimating phenotypic correlation

Both metaCCA and LD score regression has its advantages and limitations on estimating phenotypic correlation. In this section, we summarized the practical difference between the two to inform the PhenoSpD users on how to choose the appropriate methods.

MetaCCA can be applied to almost all GWASs (e.g. in our simulation, the sample size>300 and the number of SNPs>1000). However, 1) the genetic effect of SNPs may bias the phenotypic correlation estimation; 2) it only provides the central estimate of the phenotypic correlation; 3) it is difficult to quantify the effect of sample overlap.

LD score regression is designed to estimate genetic correlation between a pair of human traits. As a side product, it also provides the pair-wise phenotypic correlation estimation with standard errors. It deals with sample overlap automatically (when there is no sample overlap between two GWASs, the phenotypic correlation estimation will be zero). However, its application is limited to traits with large sample size (e.g. N > 5,000), good SNP coverage (e.g. number of SNPs > 200,000) and heritable (SNP heritability at least > 0.05) to fit the assumptions of LD score regression (Bulik-Sullivan et al., 2015a).

### The phenotypic correlations of the human metabolome

In a real case study, we applied both metaCCA and LD score regression to the human metabolome. We firstly estimated 108,978 pair-wise phenotypic correlations among these 123 metabolites from Kettunen et al (Kettunen et al, 2016) and 452 metabolites from Shin et al (Shin et al, 2014). More details of the metabolites were listed in Table S2. The phenotypic correlations estimated by metaCCA were presented in Table S5 and S6. Among the 123 metabolites from Kettunen et al, we further selected 107 of them (Table S3), which fit the assumptions of LD score regression analysis (details of the assumptions were listed in methods section). We then estimated 5,618 pair-wise correlations using LD score regression. The phenotypic correlation structure estimated by LD score regression was presented in Table S7.

### Multiple testing correction of the human metabolome

Table 2 shows the number of independent traits for two high-dimensional, complex metabolites datasets. PhenoSpD using GWAs results suggested 324.4 independent tests among 452 metabolites from Shin et al., which is close to 296 independent tests estimated using real phenotypic correlation. For metabolites from Kettunen et al, PhenoSpD suggested 44.9 number of independent tests for theses metabolites, which greatly reduced the dimensionality for these complex molecular traits. More details of the multiple testing correction are listed in Table S8.

**Table 2.**
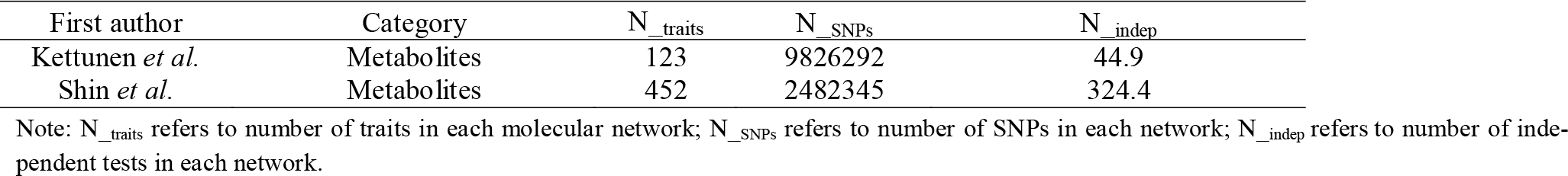
Summary of number of independent traits for the two complex metabolites networks.

## Discussion

In this study, we present an integrative method which allows phenotypic correlation estimation and multiple testing correction for human phenome using GWAS summary statistics. We illustrate the application of PhenoSpD by estimating the phenotypic correlation structure of the correlation structure of 123 metabolites from Kettunen’s study for the very first time (Kettunen et al, 2016). These results showcase the ability of PhenoSpD to estimate an appropriate multiple testing correction for complex molecular traits.

### Advantages and limitations of PhenoSpD

There are two key advantages of PhenoSpD. Firstly, PhenoSpD integrated the phenotypic correlation estimation function of metaCCA and LD score regression, with the spectral decomposition of matrices, which provides a simple way of correcting multiple testing for the human phenome using only GWAS summary results. Secondly, such multiple testing correction is still stringent (since limited sample overlap between two GWASs will drive correlated traits towards null), but more appropriate than Bonferroni correction, which is particularly valuable for GWAS of complex molecular traits. It can also be used as an indicative threshold for post-GWAS data mining tools such as MR-Base and LD Hub. For limitation, PhenoSpD can only be used for human traits from the same sample, which is a general limitation of estimating phenotypic correlation using GWAS summary statistics.

## Availability of Data and Materials

Project name: PhenoSpD

Project home page: https://github.com/MRCIEU/PhenoSpD

License: PhenoSpD is licensed under GNU GPL v3.

All data used in this manuscript are publically available and can be downloaded from the following links.

GWAS results from Shin et al:http://mips.helmholtz-muenchen.de/proj/GWAS/gwas/gwas_server/shin_et_al.metal.out.tar.gz

GWAS results from Kettunen et al:http://www.computationalmedicine.fi/data#NMR_GWAS

Operating systems: Linux, OS X, windows

Programming languages: R

### Funding

This work was supported by the Medical Research Council (MC_UU_12013/4 and MC_UU_12013/8). This work was in part supported by Cancer Research UK programme grant number C18281/A19169 (the Integrative Cancer Epidemiology Programme). P.H. is a Cancer Research UK Population Research Fellow (grant number C52724/A20138).

## References

Bulik-Sullivan, et al. (2015a) LD Score Regression Distinguishes Confounding from Polygenicity in Genome-Wide Association Studies. Nat. Genet., 47, 291–295.

Bulik-Sullivan. et al. (2015b) An atlas of genetic correlations across human diseases and traits. Nat. Genet., 47, 1236–1241.

Cichonska. et al. (2016) metaCCA: summary statistics-based multivariate meta-analysis of genome-wide association studies using canonical correlation analysis. Bioinformatics 32 (13): 1981–1989.

Hemani G, et al. (2016) MR-Base: a platform for systematic causal inference across the phenome using billions of genetic associations. bioRxiv. doi: https://doi.org/10.1101/078972

Kettunen et al. (2016) Genome-wide study for circulating metabolites identifies 62 loci and reveals novel systemic effects of LPA. Nat Commun. 7:11122.

Nyholt DR. (2004) A simple correction for multiple testing for SNPs in linkage disequilibrium with each other. Am J Hum Genet 74(4):765–769.

Li J, Ji L. (2005) Adjusting multiple testing in multilocus analyses using the eigenvalues of a correlation matrix. Heredity 95:221–227

Shin SY et al. (2014) An atlas of genetic influences on human blood metabolites. Nat Genet. 46(6):543–50.

Zheng et al. (2017) LD Hub: a centralized database and web interface to perform LD score regression that maximizes the potential of summary level GWAS data for SNP heritability and genetic correlation analysis. Bioinformatics. 33 (2): 272–279.

